# Exploring neural mechanisms underlying error-related impairments in active working memory suggests an adaptive shielding of contents during cognitive control

**DOI:** 10.1101/2025.07.19.665355

**Authors:** Yoojeong Choo, Kirsten C.S. Adam, Jan R. Wessel

**Affiliations:** Department of Psychology, University of Maryland, College Park, MD; Department of Psychological and Brain Sciences, University of Iowa, Iowa City, IA; Cognitive Control Collaborative, University of Iowa, Iowa City, IA; Department of Psychological Sciences Rice University, Houston, TX; Department of Neurology, University of Iowa Hospitals and Clinics, Iowa City, IA

## Abstract

Goal-directed behavior relies on cognitive flexibility – the ability to rapidly adapt ongoing thoughts and behaviors while preserving task-relevant information. The performance monitoring system optimizes such behavior by detecting and evaluating errors, while the working memory (WM) system maintains relevant information and protects it from interference. We investigated how these two systems interact. In prior work (Wessel et al., 2022), we found that motor errors impaired active WM maintenance (Error-Related Impairment of Active working Memory; ERIAM). Here, we aimed to identify the source of ERIAM by tracking a neurophysiological marker of visual WM maintenance – the contralateral delay activity (CDA) – throughout the error-making process. Forty-two human participants maintained visual information in WM while performing a motoric task during the delay period. Consistent with prior results, a significant ERIAM effect occurred: motor errors impaired WM performance. Critically, CDA amplitudes did not differ between motor correct and error trials before the flanker task, ruling out a general performance deficit. The CDA was also unaffected immediately after flankers, ruling out a perceptual interference explanation. Significant CDA differences only emerged after motor errors, supporting a genuinely error-related origin of the ERIAM effect. Contrary to prediction, however, CDA was more disrupted after correct responses than errors, and greater disruptions predicted a smaller ERIAM effect. These findings suggest that participants might store WM in multiple states to reduce interference from errors and that the CDA dynamics reflect these adaptive shielding strategies. These findings provide new insights into the source of error-related interference in active WM.

## Introduction

Goal-directed behavior relies on cognitive flexibility – the capacity to quickly adapt ongoing thoughts and actions while retaining critical information necessary for successful task performance. The performance monitoring system in the human brain optimizes this flexibility by continuously evaluating actions and detecting errors (Ullsperger et al., 2014), while the working memory (WM) system maintains information and protects it from interference. A key question is how performance monitoring unfolds and interacts with concurrently maintained information in WM. Traditionally, work on error processing has focused on adaptive, strategic processes that ultimately enhance subsequent behavior (Dutilh et al., 2012; Rabbitt, 1969; Ullsperger & Szymanowski, 2004). However, recent evidence suggests a contrasting perspective: errors may actually impair some ongoing processes (Ullsperger & Danielmeier, 2016; Wessel, 2018), such as perceptual attention (Beatty et al., 2018; Buzzell et al., 2017; Decker et al., 2020) and motor responses (Ceccarini et al., 2019; Guan & Wessel, 2022; Notebaert et al., 2009). This maladaptive view of error processing is further supported by recent computational modeling works, which suggest that the detrimental effects of errors seem rather inevitable (Adkins et al., 2024; Purcell & Kiani, 2016).

To further elucidate the potentially adverse effects of error processing, we recently conducted a study to examine effect of error-related impairment in active WM, termed ERIAM (Wessel et al., 2022). In a series of experiments, participants retained items in WM while performing a motoric task during the delay period. As expected, action errors committed in the motoric task impaired performance on the final WM probe. This behavioral effect was present across 7 different experiments using different motoric tasks and WM contents (Wessel et al., 2022) and has since been replicated by other groups (Brochet et al., 2025). However, behavioral measurement alone cannot answer when (or how) exactly this impairment emerged during the delay period. Here we utilized a well-validated event-related potential (ERP) component – the contralateral delay activity (CDA) – which purportedly reflects the amount of information actively maintained in WM (Adam et al., 2018; Luria et al., 2016; Vogel & Machizawa, 2004). Thusly, reductions in CDA amplitude can be interpreted as information dropping from active WM maintenance (Hakim et al., 2020). By tracking CDA, we aimed to identify the source and timing of the ERIAM effect.

In the present study, participants performed a continuous delayed estimation task while performing an arrow-flanker task during the delay period. This task was adapted from our previous work on the ERIAM effect (Wessel et al., 2022, Experiment 7), but optimized for EEG. We aimed to test three possible explanations of the ERIAM effect. First, **H1**) ERIAM may be driven by a trial-wise general performance decrement, causing mistakes on both the motor and WM part of the same trial (e.g., Cheyne et al., 2009). This hypothesis predicts that trials with motor errors may also contain subpar encoding of the WM contents, ultimately leading to errors on the probe. Second, **H2)** ERIAM could be triggered by perceptual interference from the onset of flanker task. This hypothesis predicts that on trials with motor errors, the CDA will decrease after the flanker stimuli have onset, but before a motor response has been initiated. Third, **H3)** ERIAM may be genuinely driven by error commission in the flanker task. This hypothesis predicts that the CDA will be disrupted following errors in the flanker task compared to CDA following correct responses.

Accordingly, in the present study, we measured CDA across three time windows during the delay period to test these competing explanations. Once we observed a significant difference in CDA between error and correct trials, we further examined whether this change in CDA could predict the ERIAM effect.

## Materials and Methods

### Participants

Forty-six healthy young adults participated in the experiment. Four participants were excluded from the analysis: two due to non-completion of the task; one due to chance-level accuracy (50%) in the flanker task, and one due to a technical problem during the experiment, leaving 42 participants in the analysis (mean age: 20.38 years; *SD*: 2.76; 3 left-handed, 24 females). Participants were either paid $15 per hour or received course credit for their participation. All participants had normal or corrected-to-normal visual acuity and normal color vision and did not report taking psychiatric medications. The study was approved by the ethics committee at the University of Iowa (Institutional Review Board #201511709).

Our target sample size was based on both prior empirical findings and task-specific design considerations. For detecting the behavioral ERIAM effect, an a priori power analysis using G*Power 3.1 (Faul et al., 2009) with an effect size of *d* = 0.34 – based on our previous study (Wessel et al., 2022, Exp 7) – indicated that 55 participants are required to achieve 80% power (one-tailed *t*-test, α = 0.05). For our ERP component of interest, the CDA, a simulation-based power analysis by Ngiam et al. (2021) indicated that 25 participants are sufficient to detect the component with 80% power. Given these considerations and the fact that our task design involved a high number of trials per participant and increased task difficulty to ensure a sufficient number of error trials, we targeted a sample size of approximately ∼40 participants.

### Apparatus

The experiment took place in a dimly lit experimental room. Experimental stimuli were presented on a 24-inch LCD monitor (model: BenQ XL2420B) at the native resolution of 1920 x 1080 pixels using Psychtoolbox3 (Brainard, 1997) with MATLAB 2017b (TheMathWorks, Natick, MA) run on a Linux Ubuntu computer. Participants were seated approximately 60 cm from the monitor while a chin rest was not used.

### Stimuli

#### Hybrid working memory task

The main task was designed as a hybrid of the continuous delayed estimation task (Wilken & Ma, 2004; Zhang & Luck, 2008) and the arrow-flanker task based on the paradigm from Experiment 7 in our previous study (Wessel et al., 2022). All stimuli were presented on a gray (rgb = 128 128 128) background. For the working memory task, a black fixation dot (6×6 pixels) was presented at the center of the screen. Either a left or right arrow (70×45 pixels) was used as a cue for the to-be-remembered side. Memory displays showed three colored items (60×60 pixels) on each side. Within each side, each item was vertically separated by 180 pixels. The distance from the (invisible) midline to each item was 200-212 pixels. On every trial, the locations of items were randomly jittered by ±10 pixels. Six colors were randomly chosen from a 360-color pool adopted from the previous study (Wessel et al., 2022), with the constraint that no two colors were closer than 30 degrees from each other. A color wheel (radius: 350, wheel size: 36 pixels) was presented as a WM probe. For the flanker task (Eriksen & Eriksen, 1974), five up/down arrows were used as stimuli (50×45 pixels for each).

#### Change detection task

We used a separate change detection task to measure participants’ WM capacity (Luck & Vogel, 1997; Vogel & Machizawa, 2004). We adapted the change detection task paradigm used in our previous work (Adam et al., 2017; Experimental code: https://github.com/kcsa/change-detection-task). The colors of squares were chosen from a pool of nine distinct colors: red (rgb = 255 0 0), green (0 255 0), blue (0 0 255), yellow (255 255 0), magenta (255 0 255), cyan (0 255 255), orange (255 128 0), white (255 255 255), and black (1 1 1). Stimuli were shown on a gray background. For set size 4 and 6, colors were chosen without replacement from the pool. For set size 8, colors were randomly selected from a doubled list of the colors.

### Tasks

#### Hybrid working memory task

The task paradigm is illustrated in **Figure 1**. Each trial started with a fixation dot presented at the center of the screen for 800 ms, followed by a directional arrow cue (left or right) presented above the fixation dot for 450 ms. This arrow cue indicated the to-be-remembered side. Subsequently, three colored items were presented on both the left and right sides of the hemifield for 300 ms. Participants were instructed to remember the color and the location of items only on the cued side of the display. There was a 600 ms blank delay period with a central fixation dot before the presentation of five vertically aligned arrows for the flanker task (1 central “target” arrow, surrounded by 4 flanker arrows). The five arrow stimuli were presented for 100 ms at the center. On each trial, the flanker arrows pointed in either the same direction (e.g., congruent trials) or the opposite direction (e.g., incongruent trials) as the target arrow (equal probability of congruent vs. incongruent). Participants were asked to respond to the target arrow as quickly as possible using their left hand. They used the middle and index fingers to respond to the two possible arrow directions: the key A for the up arrow and the key Z for the down arrow, respectively. Participants were asked to respond within a response time window, which was adaptively adjusted (min: 330 ms, max: 800 ms) based on their performance to ensure an adequate number of erroneous trials for each participant. If participants failed to respond within a response window, “TOO SLOW!” feedback was displayed at the center for 300 ms. The time between the flanker stimuli offset and the WM probe onset was fixed to 1500 ms. After the delay, the WM probe display was presented, featuring a color wheel with one empty black box located at one of the three color squares presented on the cued side. The color wheel was randomly rotated on each trial to discourage participants from coding colors using locations on the color wheel. Participants were instructed to report the color of probed memory item by clicking the corresponding color on the color wheel within 4000 ms. At the beginning of the WM probe, a mouse cursor was shown at the center of the screen. Once participants moved the cursor from the center of the screen, the current color at the cursor’s position was displayed as a colored square at fixation. Following the response or after the maximum time window, a blank screen with a fixation dot was presented for 800 ms and the next trial began. The experimental session consisted of 24 blocks of 32 trials, totaling 768 trials. Between blocks, participants received a summary of their performance in the previous block, including the mean accuracy (%) for the flanker task and the mean angular error for the WM probe. In addition, their cumulative performance was displayed. Cumulative mean accuracy (%) for the WM task was calculated as the percentage of WM trails where participants had an offset error of 60 degrees or less. This WM accuracy metric was solely used to encourage participants to have above 65% accuracy in both tasks. During the task, participants were instructed not to move their eyes from the fixation and to pay attention to the cued side of the items covertly. They were also asked to minimize any misses in both tasks.

**Figure 1.**
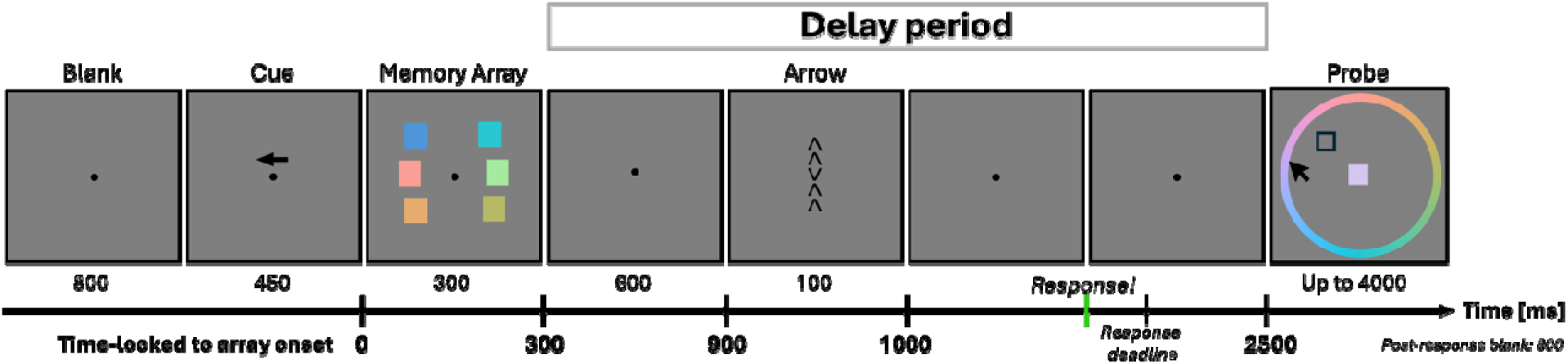
Task design. The duration of each event is shown in ms. In each trial, participants performed a continuous delayed estimation task combined with an arrow flanker task during the delay period. After the flanker task, a WM probe was presented. The response deadline for the flanker task was adjusted based on the participant’s performance while the interval between the flanker stimuli offset and the WM probe was 1500 ms. The interval from memory array onset to the WM probe was fixed as 2500 ms (Detailed timings time-locked to memory array onset were marked in bold at the bottom of the figure).

#### Change detection task

On each trial, either 4, 6, or 8 colored items were presented on the screen for 250 ms. Participants were asked to remember the color and the locations of the items. After a blank delay period of 1000 ms, a WM probe was presented at one of the remembered locations. Participants were asked to report whether the color was changed or not by pressing the “Z” key for no-change, and “/?” key for change. The probability of a change trial was 0.5. Immediately after a response, the next trial began.

### Procedure

Participants first completed 192 trials of the change detection task (96 trials of 2 blocks, 64 trials for each set size of 4, 6, and 8), followed by the main hybrid working memory task. Before starting the main task, participants underwent 1 or 2 blocks of practice, consisting of 12 trials for one block. Trials in the practice block were the same as trials in the main task, except two things: the target arrow at the center was only shown in the flanker task. In addition, participants were given feedback on responses: For a response made within a response window on the flanker task, they were given “correct” or “incorrect” feedback on the screen. For a response on the WM probe, they were provided with feedback about how far the reported color was from the original color as an angular error (e.g., “35 degrees away”). After participants fully understood the task, they moved on to the main task and completed a total of 768 trials. The total session length was up to 3 hours.

### EEG data acquisition

EEG data were recorded using a 64-channel active electrode cap connected to an actiCHamp amplifier (Brain Products) following the standard International 10-10 system. Incoming data were filtered (low cutoff = DC, high cutoff = 140 Hz) and collected at a sampling rate of 500 Hz, with channel impendence kept below 10kΩ throughout the recording session. Ground and reference channels were placed at AFz and Pz, respectively. Horizontal electrooculogram (HEOG) was not separately recorded for tracking saccadic movements; instead, saccadic artifacts were corrected using independent component analysis (ICA) during offline EEG preprocessing. Our decision was referred to previous studies reporting that ICA-based correction preserves the CDA (Drisdelle et al., 2017; Merkel et al., 2021).

### EEG preprocessing

After data acquisition, EEG data were processed using custom MATLAB scripts and the EEGLAB toolbox (Delorme and Makeig, 2004). Individual data were imported into MATLAB and downsampled to 250 Hz. Data were filtered using a least-squares finite impulse response filter (high/low cutoff: 0.1 - 30 Hz) and the continuous EEG data were epoched from -800 ms to 3000 ms relative to the cue onset (0 ms). To clean the data, we followed a two-step approach. First, non-stereotypical artifacts and noisy channels were visually inspected and removed. Rejected channels were interpolated using spherical spline interpolation, and data were re-referenced offline using an average reference. To enhance ICA decomposition, data were high-pass filtered at 1Hz before running ICA (Buzzell et al., 2017; Klug & Gramann, 2021). The filtered data were then subjected to temporal Infomax ICA (Bell & Sejnowski, 1995) with an extension to sub-Gaussian sources (Lee et al., 1999). The ICA weights were then applied to the original EEG data. Thus, all further analyses were performed on the originally filtered dataset (e.g., 0.1 Hz high-pass filtered). Next, ICs representing stereotypical artifact activities (i.e., horizontal/vertical eye movements, electrode, muscular artifacts) were visually inspected together with the automated IC classifier detection algorithm (Pion-Tonachini et al., 2019), and accordingly removed from the data. The cleaned EEG data were then used for analyses.

### Behavioral analyses

#### WM capacity assessment via change detection task

Individual WM capacity was estimated using a separate change detection task. Prior work has demonstrated that WM capacity correlates with CDA amplitude, such that individuals with higher capacity show greater CDA amplitudes (Cowan, 2001; Luck & Vogel, 2013; Luria et al., 2016; Vogel & Machizawa, 2004). Thus, it is possible that CDA differences between errors and correct responses in the flanker task, as well as the ERIAM effect in general, are affected by one’s WM capacity. Thus, accounting for WM capacity was necessary for testing the relationship between the ERIAM effect and CDA changes. In doing so, each participant’s WM capacity was computed as *K* = Set size x (Hit - False Alarm) (Cowan, 2001). K estimates for each set size (4, 6, and 8 items) were then averaged to yield one estimate per participant.

#### Hybrid working memory task

Only trials in which participants responded to both the flanker and WM tasks were included in analyses. Trials with anticipatory responses in the flanker task (RT < 150 ms) were excluded from the analysis. One participant did not understand the task until the third block, and thus these trials were not counted as valid trials in the analysis. Based on these criteria, the total exclusion rate was 10.36% of trials on average per participant (*SE* = 0.66%).

##### Flanker task

We first checked the overall error rate in the flanker task. We then examined the typical interference effects in accuracy (%) and mean response times (RT in ms), such that incongruent trials would produce poorer accuracy and slower RTs compared to congruent trials. Trials were separated as ‘motor error’ and ‘motor correct’ depending on their responses. For comparisons between error and correct conditions, only incongruent trials were included as errors in congruent trials were rare. The overall number of incongruent error trials for each participant included in the behavioral analysis were 91.62 trials (*SE* = 5.72, min: 18, max: 161). We also examined post-error processing as conduced in the previous study (Wessel et al., 2022): we examined differences in accuracy and mean RT between post-correct and post-error trials. To do so, RTs and accuracy of the next correct trials following errors and following correct were averaged respectively for each participant. We also examined the interference effect in post-error and post-correct trials to see whether errors changed performance in the next trial compared to correct responses.

##### WM task

WM performance was calculated as the average offset error for trials in the motor error and motor correct conditions. “Offset error” refers to the average angular difference (in degrees) between the original color and the reported color on a color wheel. The ERIAM was quantified by subtracting the mean offset error in motor correct trials from that in motor error trials, normalized by the mean offset error in motor correct:

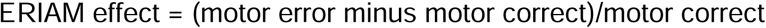

This ERIAM effect was used to examine its relationship with changes in CDA amplitudes (motor error *minus* motor correct), with only trials included in EEG analyses used for this calculation.

### EEG Analyses

Some trials were excluded from EEG analyses due to trial rejections during preprocessing. On average, each participant had 82.74 incongruent error trials available for analysis (*SE* = 5.27, min:15, max: 154). The ERIAM effect was calculated using only these trials.

#### EEG data segmentation

EEG data were segmented into two time windows: i) epochs time-locked to WM array onset (T1 epoch) and ii) epochs time-locked to the flanker response onset (T2 epoch). T1 epochs were segmented from -1200 to 2500 ms relative to WM array onset and, the T2 epochs were segmented from -1700 to 800 ms relative to the flanker response onset. ERPs for testing Hypothesis 1 – 3 were baselined relative to their respective events. For **H1**, T1 epochs were baselined using a -200 to 0 ms window relative to WM array onset. For **H2,** T2 epochs were used, but the baseline was still defined from -200 to 0 ms relative to WM array onset, since CDA was measured before the flanker response. For **H3**, T2 epochs were baselined using a window from -150 to -50 ms relative to the flanker response onset.

#### CDA analysis

To compute the CDA, we used the PO7/PO8 electrode pair, where the maximal CDA is expected based on prior work (Roy & Faubert, 2023). CDA was calculated by subtracting activity at the ipsilateral electrode from activity at the contralateral electrode, relative to the cued (to-be-remembered) side. CDA was then averaged across trials to yield one value for motor error and motor correct conditions. To test our three hypotheses, we compared CDA amplitudes between conditions across the following time windows:

For **H1**, CDA was computed from 500 to 900 ms following WM array onset. This window began 200 ms after array offset and ended at flanker task onset, capturing a period during which CDA typically stabilizes.

For **H2**, CDA was calculated from flanker task onset to flanker response. Since RTs varied by trial, the window length was trial-specific, determined by tracing backward from the flanker response to the flanker onset. Mean CDA was then computed within this window for each trial and averaged across trials separately for motor error and motor correct conditions.

For **H3**, CDA was examined following the flanker response, up to 800 ms post-response – corresponding to the final portion of the delay period (i.e., the latest possible response window). We aimed to select a window where CDA differences began to emerge and extend through the end of the trial, thus capturing mean CDA within a relevant post-response interval.

#### Brain-behavior relationships

##### i) CDA-ERIAM relationship

For each participant, the ERIAM effect was calculated as (motor error minus motor correct) / motor correct. We then tested if this behavioral effect could be predicted by the within-participant CDA difference between conditions (motor error minus motor correct) once significant CDA difference was revealed. Given that the strong association between CDA amplitude and WM capacity (Adam et al., 2018; Luria et al., 2016; Vogel & Machizawa, 2004), we computed partial correlations, controlling for individual WM capacity.

##### ii) CDA-WM capacity relationship

We examined the well-established relationship between CDA and WM capacity, which prior work has shown to be positively associated – individuals with greater WM capacity tend to show larger CDA amplitudes (i.e., greater negative deflections) (Adam et al., 2018; Luria et al., 2016; Vogel & Machizawa, 2004). This relationship was specifically evaluated within the three time windows corresponding to hypothesis **H1** – **H3**. For each window, CDA was averaged across motor error and motor correct trials to obtain a mean CDA value per participant. We then tested correlations between these values and WM capacity separately for each time window.

### Statistical Analyses

To compare CDA amplitudes between motor error and motor correct conditions, we used two-tailed paired-samples *t*-tests, with a critical p-value of 0.05. Bayes Factors were also reported for each CDA comparison to quantify the strength of evidence (Van Doorn et al., 2021). To assess relationships between CDA-ERIAM, and CDA-WM capacity, we computed Spearman rank-order correlations. For the CDA-ERIAM analysis, we additionally controlled for WM capacity using partial correlation.

## Results

### Behavioral results

#### WM capacity

Participants’ WM capacity was measured through the separate change detection task, across three set-sizes 4, 6, and 8. Participants had a mean WM capacity of 2.50 (*SE* = 0.11) [min: 1.29, max: 4.10].

#### Flanker task

Overall, participants had a 15.66% error rate in the flanker task (*SE* = 0.90%). We found the typical flanker interference effect in the flanker task. First, accuracy was significantly lower in incongruent (*M* = 72.58%, *SE* = 1.51) than congruent trials (*M* = 95.15%, *SE* = 0.58), (*t*(41) = -15.81, *p* < .001, *d* = 2.44). Second, participants made slower correct responses in incongruent (*M* = 444.59 ms, *SE* = 6.54) than congruent (*M* = 384.53 ms, *SE* = 5.41) trials, (*t*(41) = 16.06, *p* < .001, *d* = 2.478). For the following ‘motor error’ and ‘motor correct’ comparisons, only incongruent trials were included in the analyses. First, RTs in error trials were significantly faster (*M* = 356.82 ms, *SE* = 6.73) than correct trials (*M* = 444.59 ms, *SE* = 6.54) indicating that speeded responses were predominant in the motor error trials (*mean difference* = 87.77 ms, *t*(41) = 23.65, *p* < .001, *d* = 3.649). We found no difference in accuracy between post-correct (*M* = 87.02%, *SE* = 0.87) and post-error (*M* = 86.07%, *SE* = 0.94) trials, (*t*(41) = 1.44, *p* = .016, *d* = 0.221). RTs in post-error trials (*M* = 408.28 ms, *SE* = 5.36) were significantly faster than post-correct trials (*M* = 400.61 ms, *SE* = 5.92), (*t*(41) = -3.13, *p* = .003, *d* = 0.482). Post-error modulation on interference effects were not observed neither in RTs (post-correct: 57.56 ms (*SE* = 3.42) vs. post-error: 60.31 ms (*SE* = 4.18); *t*(41) = -0.92, *p* = .362, *d* = 0.142) nor in accuracy (post-correct: -19.88 ms (*SE*= 1.45) vs. post-error: -22.42 ms (*SE* = 1.45); *t*(41) = 1.65, *p* = 0.106, *d* = 0.255).

#### WM task

When only correct trials were considered, participants had greater WM errors following responses in incongruent trials (*M* = 43.74 deg, *SE* = 2.07) than congruent trials (*M* = 41.46 deg, *SE* = 2.00), (*t*(41) = 3.41, *p* = .001, *d* = 0.526). Unlike in our earlier study (Wessel et al., 2022), which used a different task design, here WM performance was affected by the congruency in the flanker task. However, no congruency effect was found for RTs. RTs for WM probe were comparable for congruent trials (*M* = 1770.37, *SE* = 37.49) and incongruent trials (*M* = 1776.96, *SE* = 38.67), (*t*(41) = -1.06, *p* = 0.294, *d* = 0.164). Next, to examine error-related impairment on WM performance (i.e., ERIAM effect), we compared WM offset errors between motor correct and motor error trials (note, this analysis was only be performed for “incongruent” trials). Participants made greater offset errors in the WM probe in the motor error (*M* = 51.66 deg, *SE* = 1.78) than motor correct (*M* = 43.74, *SE* = 2.07) trials, suggesting that error-related impairment occurred (difference = 7.92; *t*(41) = -6.29, *p* < .001, *d* = 0.97, **Figure 2A**). The ERIAM effect (*M* = 0.23, *SE* = 0.04) indicates that offset errors following motor errors were significantly increased relative to following motor correct responses (one-sample *t*-test, *t*(41) = 6.06, *p* < .001, *d* = 0.935, **Figure 2B**). Note that the mean difference in WM offset errors between conditions and ERIAM effect included in behavioral and EEG analyses were highly comparable (*r*s > 0.959, *ps* < .001). A detailed summary statistics is provided in **Appendix, Table A1**. Following errors in the flanker task, participants made faster reports in the WM probes compared to following correct responses (motor error: 1679.86 (*SE* = 38.00) vs. motor correct: 1776.96 (*SE* = 38.67), (*t*(41) = -5.49, *p* < .001, difference = -97.1 ms, *d* = 0.847).

**Figure 2.**
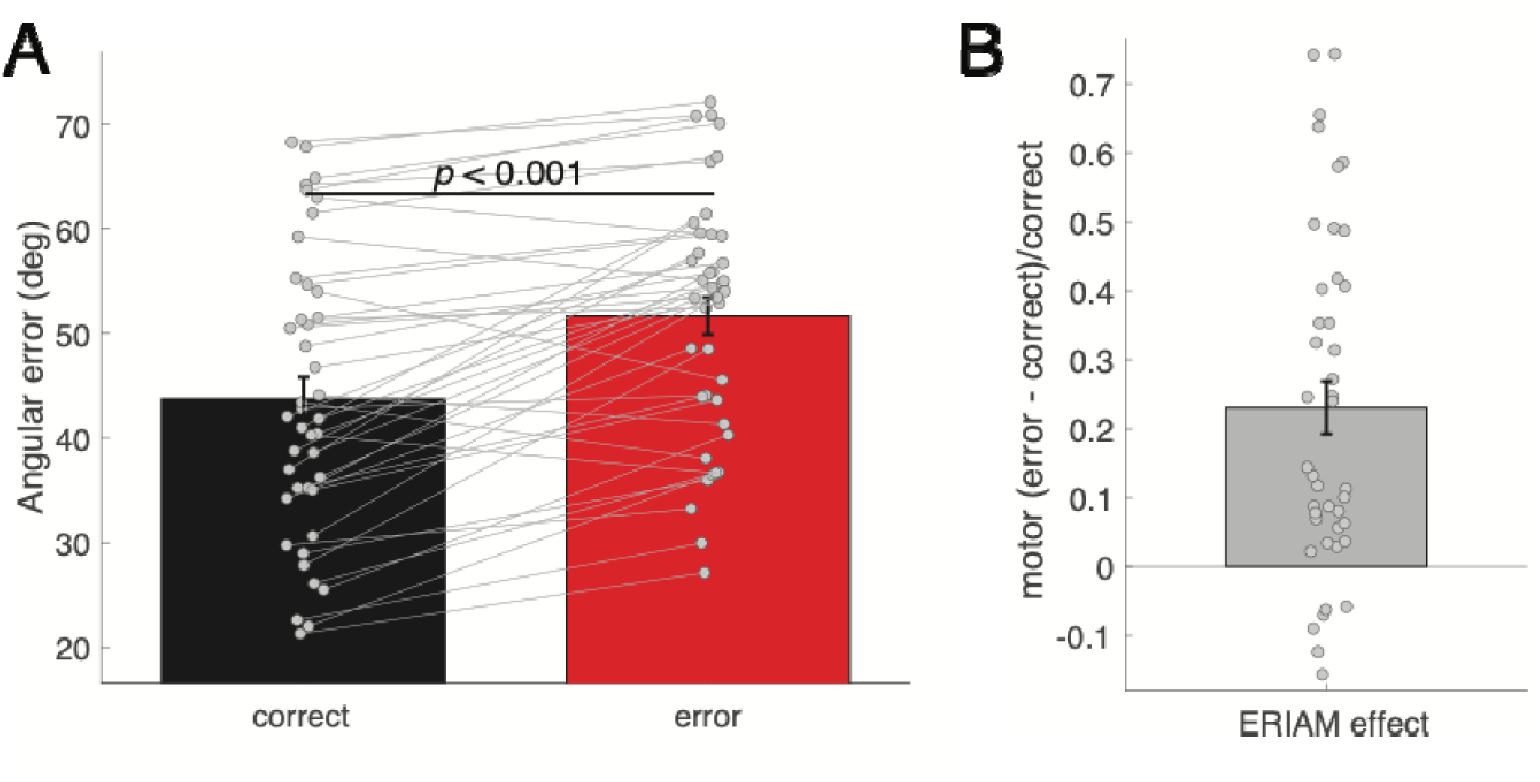
WM performance. (**A**) WM offset errors between correct (black) and error (red) in the flanker task. (**B**) The degree of ERIAM effect calculated as WM offset errors in two conditions.

### CDA Results

We analyzed differences in CDA amplitude between motor error and motor correct trials to test three hypotheses for ERIAIM effect. **H1**) general performance decrements in error trials, **H2**) differential processing of flanker task stimuli, and **H3**) error-related processes in the flanker task. To evaluate these hypotheses, we measured CDA during three time windows.

#### H1: Pre-flanker task window (500 - 900 ms after the WM array onset)

CDA was calculated from 500 to 900 ms following WM array onset (**Figure 3A**), a time window occurring after WM encoding but before the onset of the flanker task stimuli. CDA was significantly lateralized in both conditions (*ps* < .001), with no significant difference between motor error trials (*M* = -0.67, *SE* = 0.16) and motor correct trials (*M* = -0.67, *SE* = 0.15), (*t*(41) = 0.00, *p* = 0.998, *d* = 0; BF_01_ = 6.00, indicating moderate evidence for no difference). The absence of CDA difference suggests that the ERIAM effect is not attributable to generalized performance deficits or impaired encoding of WM items prior to the flanker task.

**Figure 3.**
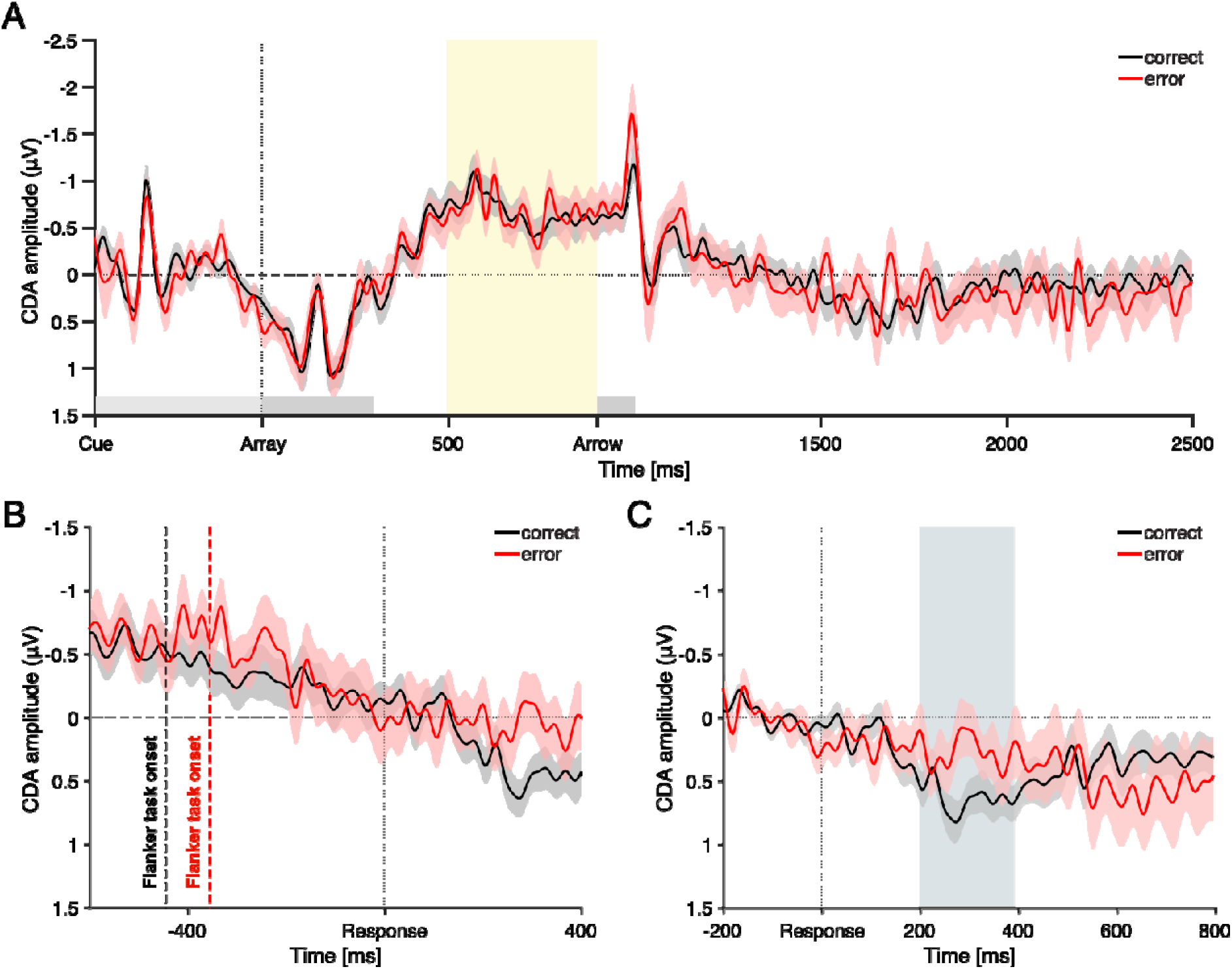
CDA dynamics across three time windows testing **H1**–**H3**. (**A**), CDA time-locked to WM array onset. The yellow shading indicates the time window of interest (500–900 ms post array) used to test **H1**. Arrow denotes flanker task onset. (**B**), CDA time-locked to the flanker response onset, used to test **H2**. Dashed lines indicate the mean flanker task onset times, traced back from response onset: -356 ms for motor error (red) and -444 ms for motor correct (black). CDA was computed for each trial between task onset and response, and then averaged across trials to yield mean CDA for each condition. (**C**), CDA time-locked to flanker response onset for testing **H3**. The gray shading marks the time window (200-396 ms post-response) where CDA differences between conditions emerged. Shaded error bars represent ± SEM.

#### H2: Post-flanker task window 1 (From flanker task onset to response)

Next, we examined whether CDA differed during the interval between the onset of the flanker task and the response (**Figure 3B**). On average, this window was longer for motor correct trials (*M* = 444 ms) than for motor error trials (*M* = 356 ms), reflecting faster erroneous responses. The CDA in the motor correct trials was significantly different from zero (*M* = -0.34, *SE* = 0.15), (*t*(41) = -2.20, *p* = 0.034, *d* = 0.339), whereas CDA in the motor error trials (*M* = - 0.41, *SE* = 0.22) did not reach significance (*t*(41) = -1.91, *p* = 0.064, *d* = 0.294). Importantly, however, there was no significant difference in CDA between the motor error and motor correct conditions, (*t*(41) = 0.43, *p* = 0.667, *d* = 0.067; BF_01_ = 5.49, indicating moderate evidence for no difference). This suggests that the ERIAM effect is unlikely to stem from early interference in the flanker task.

#### H3: Post-flanker task window 2 (After flanker response onset)

Lastly, we examined CDA amplitudes following the flanker response. Visual inspection revealed that CDA diminished in both motor error and motor correct conditions around the time of the flanker response (**Figure 3C**), showing that CDA was no longer negatively deflected. Differences between conditions began to emerge around 200 ms and persisted until 396 ms post-response (uncorrected *p* < .05; highlighted as a gray window in **Figure 3C**). CDA amplitudes were averaged within this 200 – 396 ms window for statistical analysis. In the motor error condition, CDA was not significantly different from zero (*M* = 0.27, *SE* = 0.18), (*t*(41) = 1.54, *p* = 0.132, *d* = 0.237). In contrast, the CDA in the motor correct condition was significantly greater than zero (*M* = 0.64, *SE* = 0.13), (*t*(41) = 5.04, *p* < .001, *d* = 0.778). Most notably, CDA disruption was significantly greater following correct responses than errors, (*t*(41) = 2.67, *p* = 0.011, *d* = 0.413; BF_10_ = 3.78, indicating moderate evidence for a difference). In summary, CDA was disrupted in both conditions (both rather showing positive CDA deflections), but the disruption was significantly larger following correct responses than errors – a pattern contrary to our initial predictions.

#### Relationship between CDA and the ERIAM effect

In the CDA analysis, significant amplitude differences between motor error and motor correct conditions were observed only after responses in the flanker task. We next examined whether this CDA difference could predict the behavioral ERIAM effect. Only the trials included in the EEG analyses were used to compute the ERIAM effect, which had a mean value of 0.23 (*SE* = 0.04). We assessed the relationship between CDA differences and the ERIAM effect while controlling for individual WM capacity. Results showed that greater CDA differences between motor error and motor correct trials predicted smaller ERIAM effects (partial ρ = -0.33, *p* = 0.03). It is also worth noting that the ERIAM effect was positively associated with WM capacity when controlling for CDA differences (partial ρ = 0.40, *p* = 0.009), suggesting that individual with higher WM capacity exhibited larger ERIAM effects.

#### Relationship between CDA and individual WM capacity

Lastly, we examined the association between CDA amplitude and individual WM capacity across three time windows corresponding to hypotheses **H1**–**H3**. CDA amplitudes were averaged across motor error and motor correct trials within each window. Significant negative correlations between CDA and WM capacity were observed prior to the flanker response: i) pre-flanker window (500 – 900 ms after WM array onset): (ρ = -0.53, *p* <. 001; **Figure 4A**) and ii) post-flanker task onset, pre-response window: ρ = -0.35, *p* = 0.025; **Figure 4B**). These findings replicate prior work showing that strong CDA amplitudes are associated with higher WM capacity (Vogel & Machizawa, 2004; Adam et al., 2018). However, following the flanker response, the relationship reversed, yielding a positive correlation between CDA and WM capacity (ρ = 0.33, *p* = 0.031; **Figure 4C**). This suggests that while WM capacity can be a predictor for CDA, the relationship changed to the opposite direction following motor responses, indicating that individuals with higher WM capacity exhibited greater CDA disruption following responses in general.

**Figure 4.**
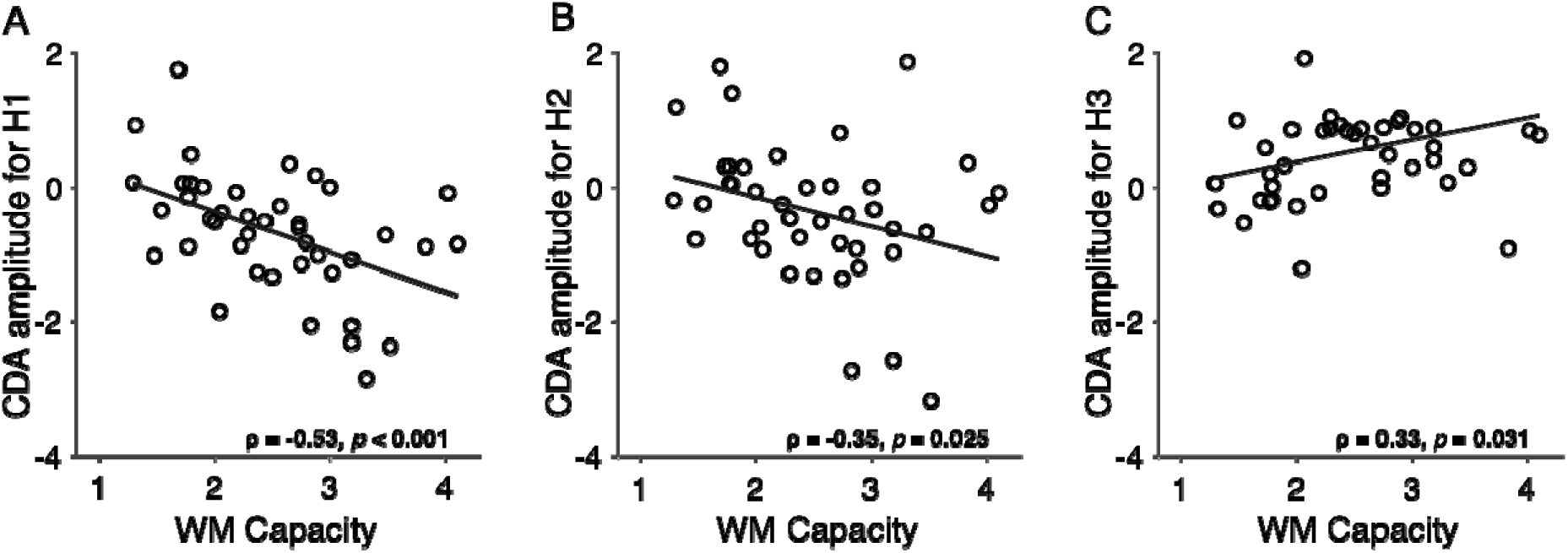
Relationship between CDA and WM capacity across three time windows corresponding to hypothesis **H1**–**H3**. (**A**) Pre-flanker task window: 500–900 ms following memory array onset (used for testing **H1**). (**B**) Response preparation window: from the flanker task onset to the flanker response (used for testing **H2**). (**C**) Post-response window: 200–396 ms following flanker response onset (used for testing **H3**). Each panel shows the correlation between CDA amplitude and individual WM capacity for the respective window.

## Discussion

In the present study, we investigated the neural source of error-related impairments in active working memory (ERIAM; Wessel et al., 2022). Using the CDA - a neural marker of active WM maintenance during delay periods (Luria et al., 2016; Vogel & Machizawa, 2004) - we aimed to determine the temporal locus of these impairments, thereby identifying the processing stage at which the ERIAM effect emerges. Our findings revealed that CDA significantly differed between motor error and correct trials only after responses in the flanker task, suggesting that motor errors directly contribute to the ERIAM effect while ruling out several alternative explanations.

### Testing three key time windows for CDA and ERIAM effect

Since the ERIAM effect could arise from distinct cognitive mechanisms, we tested CDA dynamics in three key time windows corresponding to three hypotheses:

#### **H1**: General performance decrement

The ERIAM effect could result from general lapses in WM encoding or maintenance, leading to reduced CDA (Adam et al., 2018; McCollough et al., 2007) even prior to the motor error, and possibly increasing the likelihood of making motor errors. That is, the WM deficit is not due to the motor error itself, but both are caused by a third factor. However, our findings did not support this explanation: CDA amplitudes prior to flanker task did not differ between correct and error trials (**Figure 3A**), indicating comparable WM encoding and maintenance across conditions, even when a motor error followed. Thus, general performance decrements do not explain the ERIAM effect (Adam et al., 2015; Hakim et al., 2020, 2021; Weissman et al., 2006). Supporting this interpretation, EEG markers of cognitive engagement – frontal theta and lateralized alpha power – also did not differ between conditions (**Appendix Figure A3**).

#### **H2**: Perceptual interference from flanker task

Another possibility was that flanker task stimuli interfered with WM maintenance, producing the ERIAM effect. Previous studies have shown that stimuli presented during delay periods can impair WM performance (Bae & Luck, 2019; Gresch et al., 2021; Hakim et al., 2020). Although CDA was disrupted after flanker onset, the reduction before responses did not differ between motor error and correct trials (**Figure 3B**), suggesting that perceptual interference alone cannot explain the ERIAM effect.

#### **H3**: Error commission in flanker task

Lastly, the ERIAM effect may be directly tied to motor error in the flanker task. We found that CDA differences emerged only after the motor response (**Figure 3C**), implicating error commission – not general lapses or perceptual interference – as the driver of WM impairment.

Taken together, by identifying the precise time window in which CDA begins to diverge, we provide direct evidence for the cause of the ERIAM effect. Building on prior behavioral work (Wessel et al., 2022), the current study establishes that error commission itself alters active WM maintenance, offering new insights into how cognitive control and memory systems interact.

### CDA at the time of responses to the flanker task

One key observation is that the CDA began to decrease following flanker task onset and was nearly eliminated by the time participants made their responses, regardless of accuracy. This aligns with recent studies showing that CDA is highly susceptible to interference from concurrently presented stimuli (Hakim et al., 2020, 2021; Kreither et al., 2022).

Notably, despite CDA disruption, participants demonstrated adequate WM performance, suggesting that WM representations may not rely solely on visual WM storage. Instead, they may be stored in alternative states (Lewis-Peacock et al., 2012; Myers et al., 2017; Nee & Jonides, 2013; Rose et al., 2016), and the CDA may reflect only an early state. For example, items may transition into latent or activity-silent states (Myers et al., 2017; Rose et al., 2016; Stokes, 2015; Wolff et al., 2017), or may be temporarily held as offline memory representations (Foster et al., 2024; Fukuda & Woodman, 2017; Mallett & Lewis-Peacock, 2018).

These adaptive storge mechanisms are particularly plausible in dual task scenarios, where attention must be flexibly reallocated to secondary stimuli. In our study, storing WM representations in multiple states may have allowed participants to temporarily disengage from visual WM storage and reallocate resources to process the flanker task. This could explain why CDA disruption was greater following correct responses; participants who successfully offloaded WM representations might have better engaged with the flanker task and performed well on the WM probe (Luria & Vogel, 2011).

### Interpretation of the positive CDA after a response for the flanker task

Our findings also extend theories regarding top-down modulation of CDA. Prior research shows that CDA is shaped by task goals; for example, when distractors are expected, their disruptive effect on CDA is reduced – consistent with the notion that a WM gate is closed to protect contents from interference (Chatham & Badre, 2015; Gresch et al., 2021; Hakim et al., 2021; Kreither et al., 2022; Lorenc et al., 2021). In addition, positive-going CDA shifts (i.e., CDA amplitudes rising above baseline) have been observed in response to task-relevant stimuli, interpreted as reflecting redirection of attention away from previously maintained items (Hakim et al., 2021; Kreither et al., 2022). We observed a similar positive CDA following correct responses, and the greater disruption was associated with smaller ERIAM effects.

This pattern suggests that greater CDA reduction may reflect an adaptive strategy: participants who temporarily disengaged from visual WM storage, possibly by storing items in latent states or offline representations (Foster et al., 2024; Stokes, 2015; Wolff et al., 2017), were better able to shield WM from error-related interference. Thus, the positive CDA may not be merely epiphenomenal, but rather index active cognitive strategies to preserve WM fidelity.

One tentative interpretation is that individuals who use these strategies may make more motor errors but experience less WM degradation. Indeed, those with higher flanker error rates tended to show a smaller ERIAM effect (ρ = -0.29, *p* = .062). While speculative, this tradeoff suggests that flexible WM strategies reduce interference susceptibility but may impair concurrent task performance. These explanations remain post-hoc and require direct testing.

### Alternative explanations for positive CDA

While we interpret the positive CDA as reflecting adaptive control strategies, alternative explanations warrant consideration. First, prior studies show that CDA amplitude naturally decreases over time (Adam et al., 2018; Hakim et al., 2020; McCollough et al., 2007), suggesting that the observed positivity could reflect a temporal drift. However, in our data, CDA dropped rapidly after the motor response, particularly following correct responses. Importantly, a simple drift would not explain the observed relationship between CDA and the ERIAM effect, making this account less plausible.

A second possibility is that the observed positivity reflects the late directing attention positivity (LDAP) – a component associated with orienting or anticipation of an upcoming target at the cued side (Eimer et al., 2002; McDonald & Green, 2008; Nobre et al., 2000). Under this interpretation, positive CDA could reflect anticipation of the WM probe triggered by the motor response. However, it is unclear how a motor response could act as such a cue well before the WM probe appears.

Crucially, CDA amplitudes before the flanker task strongly correlated with WM capacity (**Figure 4A & 4B**) replicating prior work (Luria et al., 2016; Vogel & Machizawa, 2004). After the flanker response, however, this relationship reversed (**Figure 4C**), suggesting that the positive CDA may reflect a complex interaction between active WM maintenance and error-related processes. Future research is needed to elucidate the neural mechanisms underlying this phenomenon.

### Future research questions

Our findings raise several avenues for future investigation. First, the current study employed a fixed, short delay between the motor response and the WM probe. However, it is possible that participants re-engage visual WM storage after completing the flanker task if the delay period is extended (Clapp et al., 2010; McCollough et al., 2007). If so, CDA reactivation might be greater following correct responses than errors. Alternatively, if WM had been reconfigured into activity-silent or alternative storage states (Lewis-Peacock et al., 2012; Myers et al., 2017), CDA might not recover, and condition differences would remain minimal.

Another question concerns the temporal dynamics of error processing (Wessel, 2018). According to this framework, errors trigger automatic inhibition-attentional orienting, followed by strategic adjustments (Choo et al., 2023; Guan & Wessel, 2022; King et al., 2010). An open question is whether the ERIAM effect can be offset by manipulating the timing of post-error WM reactivation. For example, introducing a secondary WM array shortly after the motor error might be ineffective due to lingering error-related inhibition-orienting processes (Jentzsch & Dudschig, 2009; Notebaert et al., 2009). However, allowing recovery time may enable task-set updating and successful WM reinstatement, potentially reducing ERIAM effect or even improving WM. These avenues can illuminate how cognitive control supports WM under error-related interference.

## Conclusion

This study investigated the source of the ERIAM effect (Brochet et al., 2025; Wessel et al., 2022). By tracking CDA, we demonstrated that the ERIAM effect is primarily driven by error commission, rather than general task disengagement or interference from flanker task. Contrary to our original prediction, CDA disruption was greater following correct responses, and this was linked to smaller ERIAM effects. This might suggest that disengaging from visual WM storage protects memory from interference, reflecting an adaptive shielding strategy. However, this interpretation remains speculative. Future research should test this hypothesis using a priori designs aimed at dissociating multi-state storage mechanisms and their role in error-related cognitive control.

## Appendix

**Table A1.** Summary statistics of WM performance. Statistics reported are group means and SEMs between parentheses. For analyses for comparisons between motor error and motor correct conditions, only incongruent trials were included.

| WM task | Behavioral data | EEG data |
| --- | --- | --- |
| motor correct | 43.74 (2.07) | 43.67 (2.10) |
| motor error | 51.66 (1.78) | 51.37 (1.73) |
| Difference in offset errors | 7.92 (1.26) | 7.70 (1.42) |
| ERIAM effect | 0.23 (0.04) | 0.23 (0.04) |

### Exploratory EEG analyses

We conducted several exploratory analyses to further characterize the neural mechanisms underlying ERIAM effect. These analyses focused on (1) ERPs evoked by the flanker task stimuli and responses, and (2) oscillatory power changes during the T1 and T2 epochs used in the CDA analysis.

### ERP analyses

#### i) Early sensory processing: P1 and N1 ERPs

We examined whether early visual processing of the flanker stimuli differed between motor error and motor correct conditions. Specifically, we analyzed two classic ERP components: P1 reflecting initial sensory encoding and N1 associated with attentional selection; (Hillyard & Anllo-Vento, 1998; Vogel & Luck, 2000). Because flanker stimuli were centrally presented, we refer to these as global P1 and global N1. EEG data were averaged at electrodes PO7/PO8. Peaks were identified across both conditions: P1 was identified at 56 ms post-stimulus within a 50-130 ms window. N1 was identified at 156 ms within 100 – 180 ms window. Mean amplitudes were calculated using ±20 ms time windows around each peak and compared between conditions.

#### ii) Response-locked error-related ERPs: Error-Related Negativity (ERN) and error Positivity (Pe)

We also analyzed well-established ERP markers of error processing: the ERNA and Pe (Falkenstein et al., 1991; Gehring et al., 1993). These were extracted using a difference wave (motor error minus motor correct) time-locked to the response (Buzzell et al., 2017). The ERN was identified at 32 ms at the electrode FCz. Amplitudes were calculated within a ±20 ms window. The Pe was identified at 356 ms at the electrode Pz. Amplitudes were calculated within a ±100 ms window.

### EEG spectral power analysis

We examined two established EEG spectral markers: i) frontal theta (4-7Hz), and ii) lateralized alpha power (8-12 Hz), within two time windows: one prior to the flanker task and one following the flanker response. Frontal theta is considered a neural signature of cognitive control (Cavanagh & Frank, 2014), and is associated with successful WM performance (Adam et al., 2015, 2018; Itthipuripat et al., 2013). In addition, lateralized alpha power suppression reflects the deployment of covert spatial attention (e.g., Thut et al., 2006; Worden et al., 2000).

#### Signal processing and feature extraction

EEG data were bandpass-filtered using a two-way, least-squares finite impulse response filter, implemented within the built-in *eegfilt* function in EEGLAB. The Hilbert transform (hilbert.m) was then applied to extract instantaneous power values from the bandpass-filtered signals. Power was normalized by computing the percent change relative to a baseline period 300 ms before the onset of the cue (i.e., -750 to -450 ms relative to WM array onset). Frontal theta power was computed by averaging across electrodes FC1, FC2, and FCz. Lateralized alpha power was calculated from electrodes O1/O2, P7/P8, PO3/PO4, and PO7/PO8. Power was defined as the difference in percent change between contralateral and ipsilateral electrodes relative to the to-be-remembered side.

#### Time windows and statistical comparisons

Two time windows identical to those used in the CDA analysis were selected: 1) T1 epoch (500 – 900 ms post-WM array onset) to assess attentional and control-related dynamics prior to the flanker task. 2) post-response window (-200 to 800 ms relative to flanker response) to examine potential changes in oscillatory power driven by error-related processes. Power differences between motor error and motor correct trials were compared at each time point within each window. To correct for multiple comparisons, we applied the False Discovery Rate (FDR) correction procedure (Benjamini et al., 2006).

### Results

#### Sensory processing: P1 and N1 ERPs

P1 and N1 components evoked by the flanker task stimuli are shown in Figure A1. The P1 component, measured within a 76 – 116 ms window, did not differ significantly between motor error (*M* = 4.54, *SE* = 0.61) and motor correct (*M* = 4.66, *SE* = 0.58) conditions, (*t*(41) = -0.88, *p* = 0.385, *d* = 0.136). In contrast, the N1 component, measured within a 136 – 176 ms window, was significantly larger in motor error (*M* = -6.82, *SE* = 0.92) than motor correct (*M* = -6.47, *SE* = 0.92) conditions, (*t*(41) = -2.04, *p* = 0.048, *d* = 0.314) while the effect size was small. Together, these findings indicate that early perceptual encoding (P1) was similar across conditions, whereas attentional selection (N1) was heightened in error trials. This enhanced processing was not captured by the main CDA analysis (**Figure 3B**), suggesting that flanker-related processing differences may be reflected in transient ERPs rather than sustained WM-related activity.

**Figure A1.**
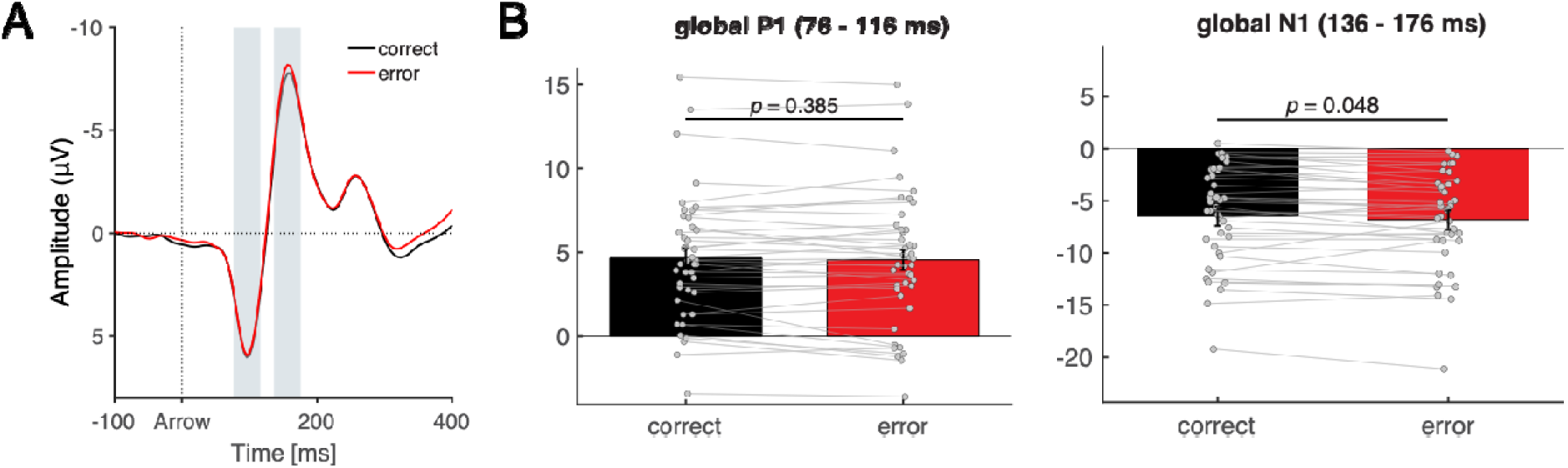
Early sensory processing of flanker task stimuli. **(A)** Time windows used for calculating P1 (76 – 116 ms) and N1 (136 – 176 ms) are shown as gray shaded areas. Shaded error bars represent ± SEM. (**B**) Mean P1 and N1 amplitudes in motor correct (denoted “correct”) and motor error (denoted “error”) conditions.

#### Error-related ERPs: ERN and Pe

We examined two well-established error-related ERP components: the ERN and the Pe. The difference waveforms (motor error minus motor correct) for each component are shown as green lines in **Figure A2**, along with their corresponding scalp topographies below. As expected, both components were significantly enhanced following errors. The ERN was significantly more negative following errors (*M* = -2.88, *SE* = 0.39) than correct responses (*M* = 1.22, *SE* = 0.34), (*t*(41) = -10.22, *p* < .001, *d* = 1.577). The Pe was significantly more positive following errors (*M* = 1.91, *SE* = 0.51) than correct responses (*M* = -1.14, *SE* = 0.46), (*t*(41) = 7.87, *p* < .001, *d* = 1.214). These robust effects confirm the presence of typical error-related ERPs, consistent with prior literature on performance monitoring.

**Figure A2.**
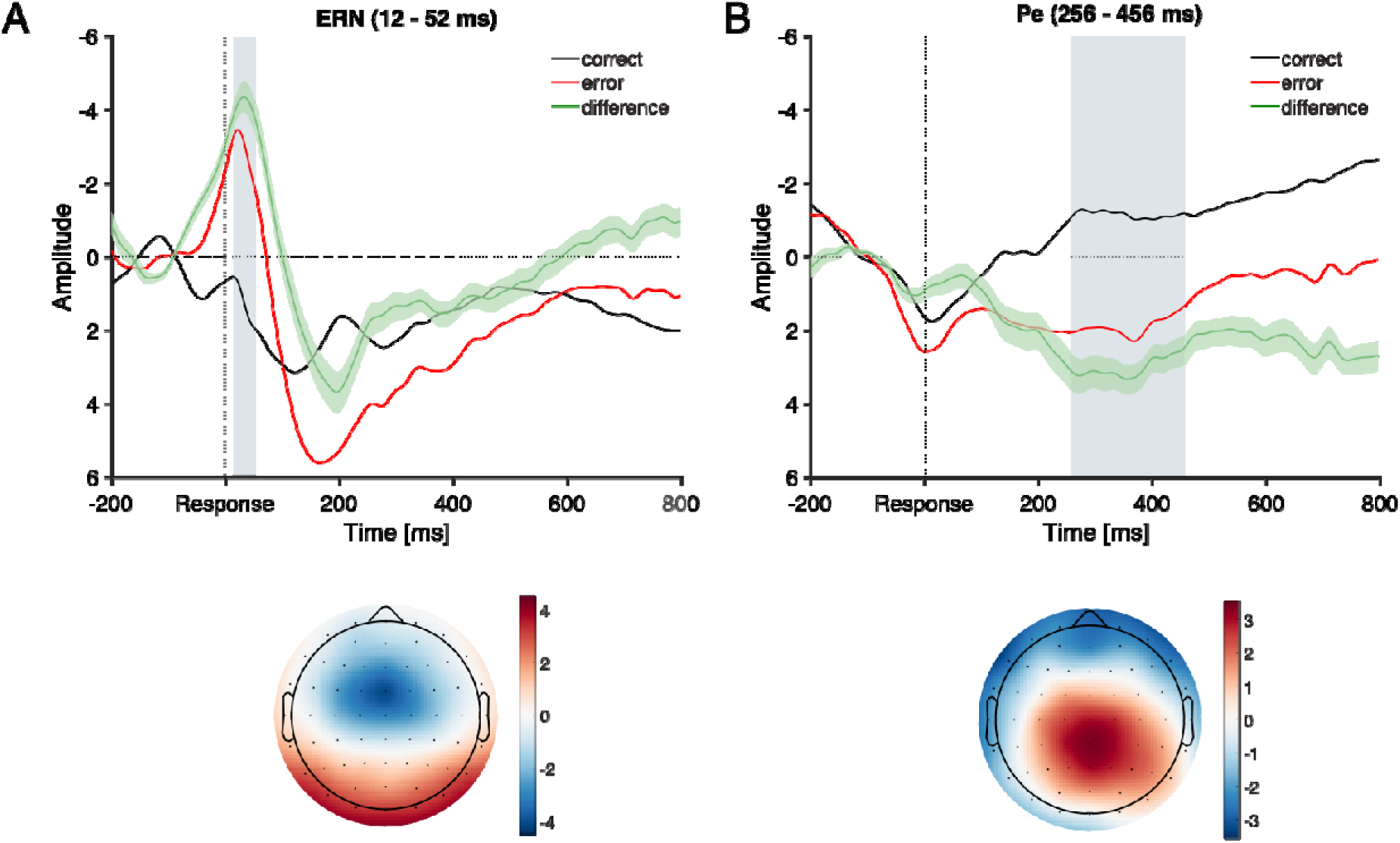
Error-related ERPs. (**A**) The ERN measured at FCz (top), with its corresponding topography shown below. The gray shaded window indicates the analysis window surrounding ERN peak (12 – 52 ms). (**B**) The Pe measured at Pz (top), and its topography shown below. The gray shaded window indicates the Pe analysis window (256 – 456 ms). Shaded error bars represent ± SEM.

#### Lateralized alpha power

We tested whether a sustained covert spatial attention toward the cued side could account for the ERIAM effect. To this end, we analyzed lateralized alpha power suppression, a well-established index of sustained attention to one visual hemifield, across both conditions at two key time windows (**Figure A3-A**). In the 500 – 900 ms window following WM array onset, lateralized alpha power did not significantly differ between motor error (*M* = -9.42, *SE* = 8.80) and motor correct (*M* = -12.17, *SE* = 5.54) conditions, (*t*(41) = 0.32, *p* = 0.75, *d* = 0.05). Similarly, in the -200 to 800 ms window (time-locked to flanker response onset), no significant difference was observed between conditions. These findings suggest that participants allocated and sustained covert spatial attention to the to-be-remembered side to a similar extent, regardless of whether their flanker response was correct or erroneous.

#### Frontal theta power

We tested whether differences in cognitive control engagement between motor correct and motor error trials could account for the ERIAM effect. To this end, frontal theta power was measured during two key time windows **(Figure A3-B)**. In the 500 – 900 ms window following WM array onset, frontal theta power did not significantly differ between motor error (*M* = 107.85, *SE* = 11.24) and motor correct (*M* = 100.46, *SE* = 6.02) conditions, (*t*(41) = -0.75, *p* = 0.457, *d* =0.116), suggesting that the cognitive demands prior to the flanker task were comparable across conditions. In contrast, during the window surrounding the flanker response (-56 – 384 ms, FDR-corrected *p* < .05, time-locked to response onset), frontal theta power was significantly increased for motor error trials (*M* = 322.29, *SE* = 31.39) than motor correct trials (*M* = 165.86 *SE* = 8.64), (*t*(41) = 5.73, *p* < .001, *d* = 0.884). This reflects a typical error-related increase in frontal theta activity, indicating heightened cognitive control demand following erroneous responses (Cavanagh & Frank, 2014).

**Figure A3.**
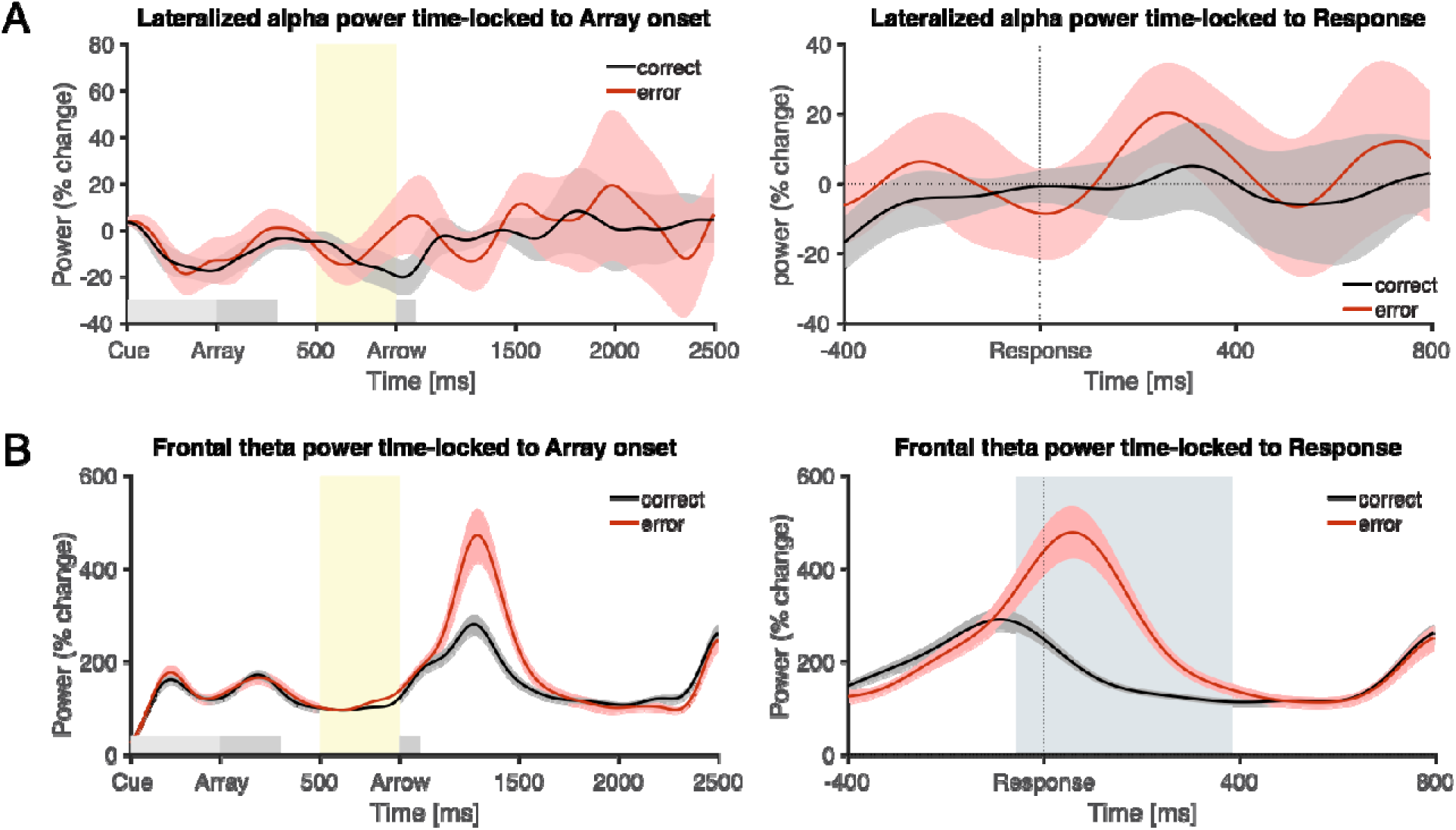
Spectral power analysis. (A) Lateralized alpha power time-locked to WM array onset (left) and time-locked to flanker task response (right). (B) Frontal theta power time-locked to WM array onset (left) and time-locked to flanker task response (right). Yellow highlighting denotes the time window of interest. Grey highlighting indicates the window testing difference between two conditions time-locked to Response. Shaded error bars represent ± SEM.

## Acknowledgements

This work was supported by National Institute of Health (R01NS102201) to JRW. We would like to thank Nathan Chalkley for help with data collection. We also thank the Human Perception and Performance Group (in particular, Andrew Hollingworth, Cathleen Moore, Toby Mordkoff, and Jiefeng Jiang) at the University of Iowa for helpful feedback and suggestions.

## Data Availability Statement

Datasets as well as related analysis scripts will be made publicly available upon acceptance of the manuscript at https://osf.io/sjuy7/.

## Diversity in Citation Practices

Retrospective analysis of the citations in every article published in this journal from 2010 to 2021 reveals a persistent pattern of gender imbalance: Although the proportions of authorship teams (categorized by estimated gender identification of first author/last author) publishing in the *Journal of Cognitive Neuroscience* (*JoCN*) during this period were M(an)/M = .407, W(oman)/M = .32, M/W = .115, and W/W = .159, the comparable proportions for the articles that these authorship teams cited were M/M = .549, W/M = .257, M/W = .109, and W/W = .085 (Postle and Fulvio, *JoCN*, 34:1, pp. 1-3). Consequently, *JoCN* encourages all authors to consider gender balance explicitly when selecting which articles to cite and gives them the opportunity to report their article’s gender citation balance. The authors of this paper report its proportions of citations by gender category to be: M/M = .721; W/M = .167; M/W = .075; W/W = .046.

## Author Contributions (CRediT)

Yoojeong Choo, Conceptualization, Data curation, Formal analysis, Investigation, Methodology, Software, Visualization, Writing - original draft, Writing - review and editing

Kirsten C.S Adam, Methodology, Supervision, Writing - review and editing

Jan R. Wessel, Conceptualization, Methodology, Project administration, Resources, Funding acquisition, Supervision, Writing - review and editing;

